# A *trans*-complementation system for SARS-CoV-2

**DOI:** 10.1101/2021.01.16.426970

**Authors:** Xianwen Zhang, Yang Liu, Jianying Liu, Adam L. Bailey, Kenneth S. Plante, Jessica A. Plante, Jing Zou, Hongjie Xia, Nathen Bopp, Patricia Aguilar, Ping Ren, Vineet D. Menachery, Michael S. Diamond, Scott C. Weaver, Xuping Xie, Pei-Yong Shi

## Abstract

The biosafety level-3 (BSL-3) requirement to culture severe acute respiratory syndrome coronavirus 2 (SARS-CoV-2) is a bottleneck for research and countermeasure development. Here we report a *trans*-complementation system that produces single-round infectious SARS-CoV-2 that recapitulates authentic viral replication. We demonstrate that the single-round infectious SARS-CoV-2 can be used at BSL-2 laboratories for high-throughput neutralization and antiviral testing. The *trans*-complementation system consists of two components: a genomic viral RNA containing a deletion of ORF3 and envelope gene, and a producer cell line expressing the two deleted genes. *Trans-*complementation of the two components generates virions that can infect naive cells for only one round, but does not produce wild-type SARS-CoV-2. Hamsters and K18-hACE2 transgenic mice inoculated with the complementation-derived virions exhibited no detectable disease, even after intracranial inoculation with the highest possible dose. The results suggest that the *trans*-complementation platform can be safely used at BSL-2 laboratories for research and countermeasure development.

## INTRODUCTION

Three zoonotic betacoronaviruses have emerged to cause global epidemics or pandemics in less than twenty years: severe acute respiratory syndrome coronavirus (SARS-CoV) in 2002, Middle East respiratory syndrome coronavirus (MERS-CoV) in 2012, and SARS-CoV-2 in 2019 (*1*). The coronavirus disease 2019 (COVID-19) pandemic has caused unprecedented social and economic disruption. As of January 16, 2021, SARS-CoV-2 had infected over 94 million people, leading to over 2 million deaths (https://www.worldometers.info/coronavirus/). In response to the pandemic, the scientific community has rapidly developed experimental platforms to study COVID-19 and to develop countermeasures. Several groups have established infectious cDNA clones and reporter SARS-CoV-2 to facilitate the development and analysis of first-generation vaccines and therapeutics (*2-6*). However, since SARS-CoV-2 is a biosafety level-3 (BSL-3) pathogen, the requirement of high containment represents a bottleneck for antiviral and vaccine evaluation. Thus, a BSL-2 cell culture system that recapitulates authentic viral replication is urgently needed.

The genome of SARS-CoV-2 is a positive-sense, single-stranded RNA of approximately 30 kb in length. The SARS-CoV-2 virion consists of an internal nucleocapsid [formed by the genomic RNA coated with nucleocapsid (N) proteins] and an external envelope [formed by a cell-derived bilipid membrane embedded with spike (S), membrane (M), and envelope (E) proteins] (*7*). The genomic RNA encodes open-reading-frames (ORFs) for replicase (ORF1a/ORF1b), S, E, M, and N proteins, as well as seven additional ORFs for accessory proteins (1). Stable cell lines containing replicons (self-replicating viral RNA genomes with one or more gene deletions) have been developed for many viruses, including coronaviruses (*8-12*). Because replicons lack structural genes, they are not infectious and can safely be manipulated in BSL-2 laboratories. For SARS-CoV-2, although a transient replicon system has been established (*13*), no stable replicon cell line has been reported. To overcome this gap, we have developed a single-round infectious SARS-CoV-2 through *trans-*complementation. The single-round SARS-CoV-2 is engineered with a reporter gene that facilitates high-throughput antiviral screening and neutralizing antibody measurement. We validated the safety of the system in cell cultures, hamsters, and highly susceptible human angiotensin-converting enzyme 2 (hACE2) transgenic mice. Our results suggest that the *trans-*complementation system can be used safely at BSL-2 laboratories.

## RESULTS

### A single-round infectious SARS-CoV-2 system

**Fig. 1A** depicts the *trans*-complementation system to produce single-round infectious SARS-CoV-2. The system contains two components: (i) a viral RNA containing a mNeonGreen (mNG) reporter gene and a deletion of ORF3 and E genes (ΔORF3-E; **Fig. 1B**) and (ii) a Vero E6 cell line expressing the ORF3 and E proteins under a doxycycline inducible promoter (Vero-ORF3-E; **Fig. 1C-D**). Upon electroporation of ΔORF3-E RNA into Vero-ORF3-E cells and addition of doxycycline, *trans-* complementation enables production of virions that can continuously infect and amplify on Vero-ORF3-E cells; however, these virions can only infect normal cells for a single round due to the lack of ORF3 and E genes in the packaged RNA genome (**Fig. 1A**).

**Figure 1.**
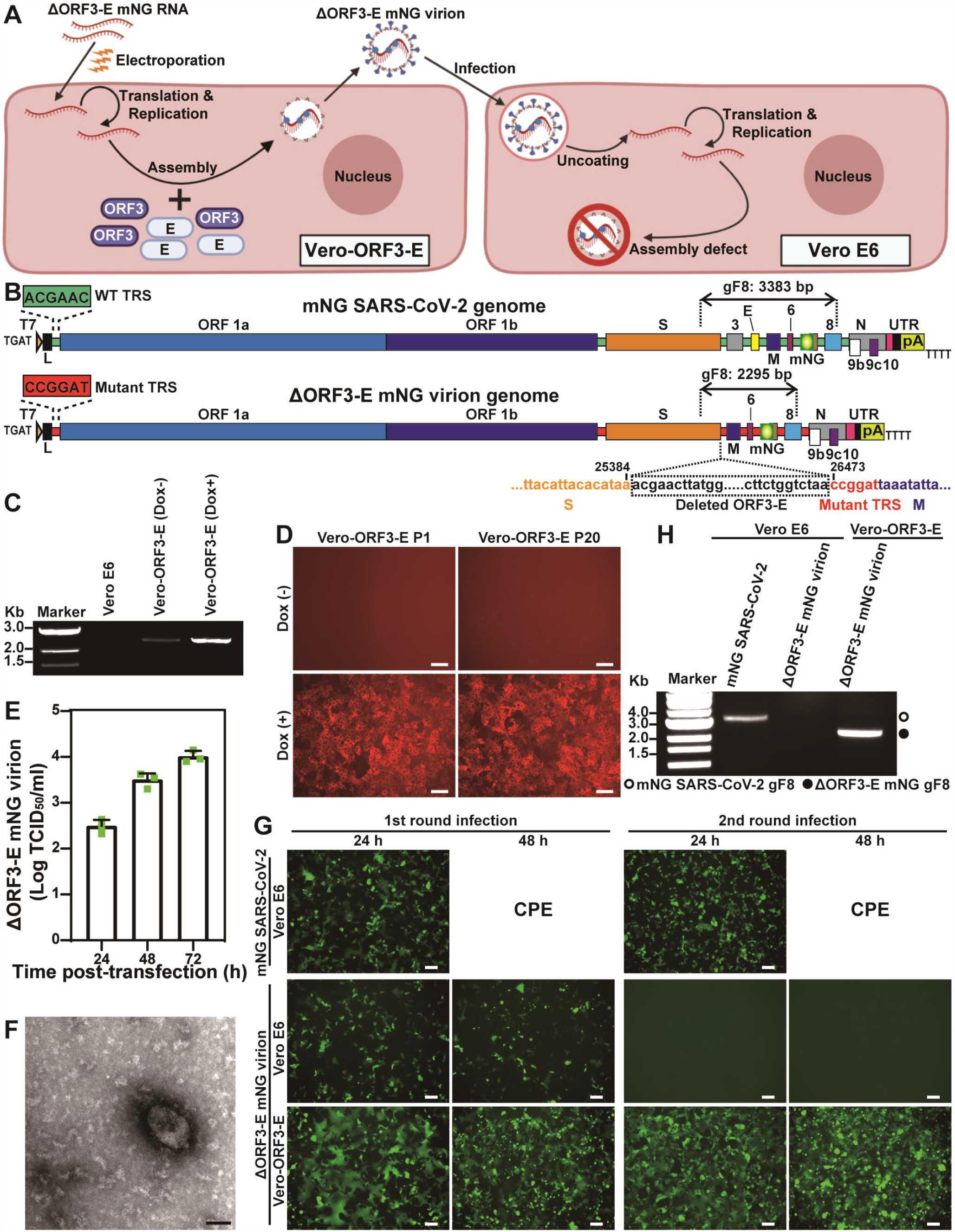
Generation of single-round infectious ΔORF3-E mNG virion. (**A**) A *trans*-complementation system for SARS-CoV-2. Vero-ORF3-E cells are electroporated with ΔORF3-E mNG RNA. *Trans-*complementation produces ΔORF3-E mNG virion (*left panel*) which can infect naïve Vero E6 cells for only single round (*right panel*). (**B**) ΔORF3-E mNG virion genome. Both the full-length mNG SARS-CoV-2 genome (*top panel*) and the ΔORF3-E mNG virion genome (*bottom panel*) are shown. The genomic fragment 8 (gF8) of RT-PCR analysis are indicated above both genomes. The ORF3-E deletion junction is indicated. The WT and mutant Transcription Regulatory Sequences (TRS) are also depicted. (**C**) ORF3-E RNA expression in Vero-ORF3-E cells. Doxycycline (Dox) was used to induce the expression of ORF3-E RNA. RT-PCR analyses were performed on Vero-ORF3-E cells with or without doxycycline induction as well as on naïve Vero E6 cells. (**D**) Induction of mCherry expression in Vero-ORF3-E cells. Passage 1 (P1) and 20 (P20) of Vero-ORF3-E cells were induced by doxycycline to express mCherry fluorescence. Scale bar, 100 μm. (**E**) Production of ΔORF3-E mNG virion after electroporation. After electroporating ΔORF3-E mNG RNA into Vero-ORF3-E cells (with doxycycline), infectious titers of ΔORF3-E mNG virion were measured from culture medium. Three sets of repeated experiments are presented with bars representing standard deviations. (**F**) Negative-staining electron microscopic image of ΔORF3-E mNG virion. Scale bar, 50 nm. (**G**) Analysis of ΔORF3-E mNG virion infection. Vero E6 or Vero-ORF3-E cells were incubated with WT mNG SARS-CoV-2 or ΔORF3-E mNG virion for 2 h. The cells were washed three times with PBS to remove residual input virus. At 48 h post-infection, the supernatants of the infected cells were transferred to fresh Vero E6 or Vero-ORF3-E cells for a second round of infection. The mNG signals from both rounds of infected cells are presented. Scale bar, 100 μm. (**H**) RT-PCR analysis. Extracellular RNA from the second-round infection from (**G**) was harvested at 48 h post-infection. Fragment 8 of the viral genome, depicted in (**B**), was amplified by RT-PCR to confirm the ORF3-E deletion and mNG retention.

Our *trans*-complementation system is engineered with several safeguards to eliminate wild-type (WT) SARS-CoV-2 production. Besides the ORF3-E deletion, the ΔORF3-E viral RNA contained two additional modifications. (i) The transcription regulatory sequence (TRS) of ΔORF3-E RNA was mutated from the WT ACGAAC to CCGGAT (mutant nucleotides underlined; **Fig. 1B**). Recombination between the TRS-mutated ΔORF3-E RNA with inadvertently contaminating viral RNA would therefore not produce replicative virus (*14, 15*). (ii) An mNG gene was engineered at ORF7 of ΔORF3-E RNA to facilitate the detection of viral replication (**Fig. 1B**). The *trans-*complementing Vero-ORF3-E cell lines were produced by transducing Vero E6 cells with a lentivirus encoding the following elements (**Fig. S1A**): a TRE3GS promoter that allows doxycycline to induce ORF3 and E protein expression (**Fig. 1C-D and S1B**); an mCherry gene that facilitates selection of cell lines with high levels of protein expression (**Fig. S1C**); a foot-and-mouth disease virus 2A (FMDV 2A) autocleavage site that enables translation of individual mCherry and viral E protein; and an encephalomyocarditis virus internal ribosomal entry site (EMCV IRES) that bicistronically translates the ORF3 protein. The above design eliminated overlapping sequences between the ORF3-E mRNA and ΔORF3-E viral RNA, thus minimizing homologous recombination during *trans-*complementation. The Vero-ORF3-E cell line stably expressed the engineered proteins after 20 rounds of passaging, as indicated by the mCherry reporter (**Fig. 1D**).

Electroporation of ΔORF3-E mNG RNA into doxycycline-induced Vero-ORF3-E cells produced virions of ~10^4^ median Tissue Culture Infectious Dose (TCID_50_)/ml titer (**Fig. 1E**). The ΔORF3-E mNG virion exhibited a diameter of ~91 nm under negative staining electron microscopy (**Fig. 1F**). The ΔORF3-E mNG virion produced in the supernatant could infect Vero-ORF3-E cells for multiple rounds, but for only one round on naïve Vero E6 (**Fig. 1G-H**), Calu-3, or hACE2-expressing A549 cells (A549-hACE2; **Fig. S2**). As controls, WT mNG SARS-CoV-2 could infect cells for multiple rounds (**Fig. S2**). These results indicate that the *trans*-complementation system produces virions that can only infect WT cells for single round.

### Adaptive mutations to improve virion production

To improve the efficiency of the *trans-*complementation platform, we serially propagated ΔORF3-E mNG virions on Vero-ORF3-E cells for 10 passages (3-4 days per passage) to select for adaptive mutations. The P10 virion replicated to higher titers than the P1 virion on Vero-ORF3-E cells (**Fig. 2A**), retained the mNG reporter (**Fig. 2B-C**), and still infected normal Vero cells for only single round (**Fig. S3**). Whole genome sequencing of the P10 virion revealed three mutations in the nsp1, nsp4, and S genes (**Fig. 2D**). Engineering of these mutations into ΔORF3-E mNG RNA showed that all three were required to enhance the *trans-*complementation efficiency, producing 10^6^ TCID_50_/ml of virions (**Fig. 2A-B**). These results indicate that (i) adaptive mutations can be selected to improve the yield of single-round virions and (ii) WT virus is not produced from the *trans-*complementation system.

**Figure 2.**
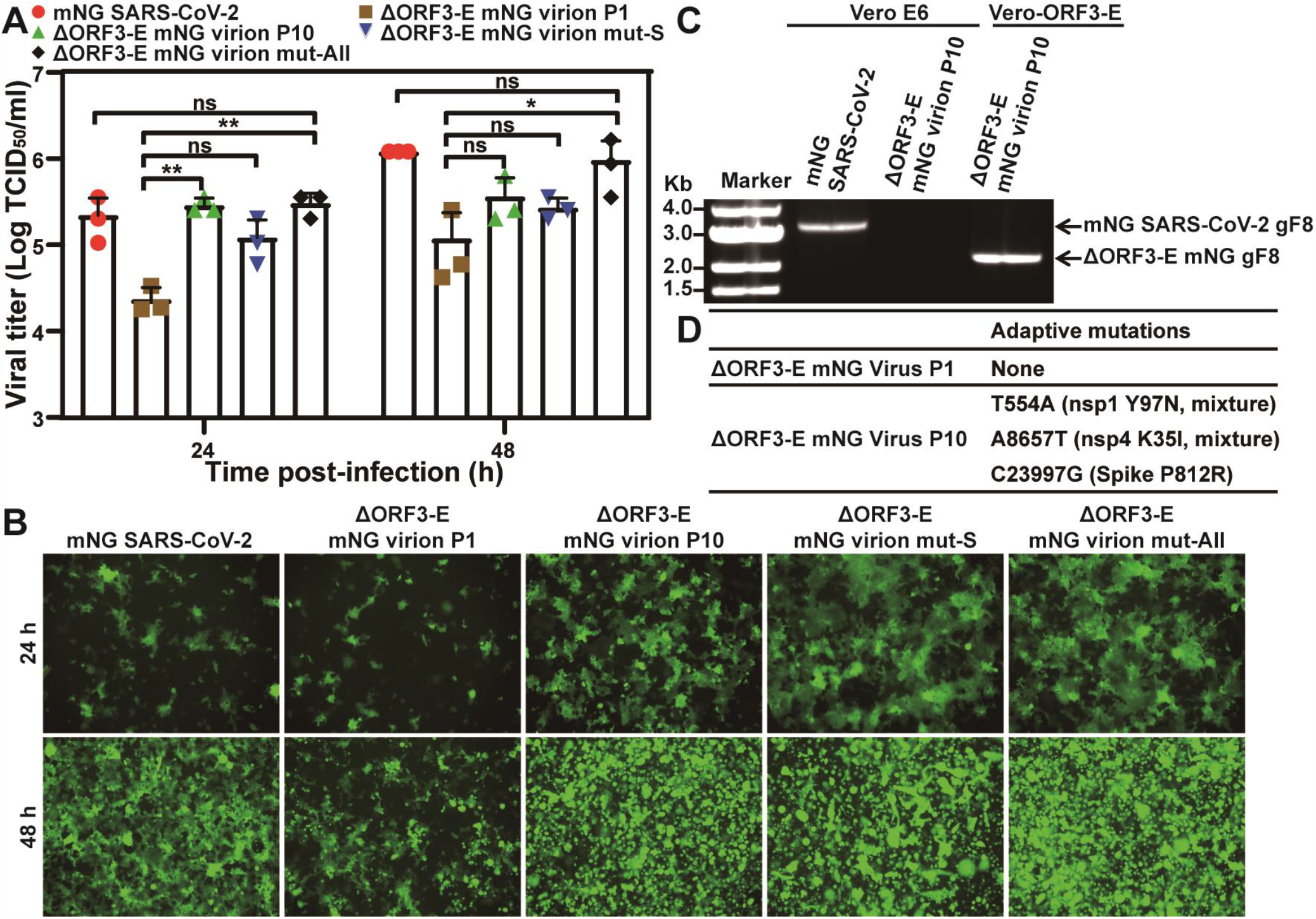
Adaptive mutations to improve the yield of ΔORF3-E mNG virion production. (**A**) Viral replication kinetics on Vero-ORF3-E cells. Adaptive mutations (**D**) were selected by continuously passaging the ΔORF3-E virion on Vero-ORF3-E cells for 10 rounds. For comparing the replication kinetics of the passaged viruses, Vero-ORF3-E cells were infected with the P1 or P10 ΔORF3-E virion, ΔORF3-E virion containing an S mutation in (**D**) [ΔORF3-E virion mut-S], or ΔORF3-E virion containing all adaptive mutations in nsp1, nsp4, and S in (**D**) [ΔORF3-E virion mut-All] at an MOI of 0.15. WT mNG SARS-CoV-2 was included as a control. Viral titers in culture supernatants are presented. ANOVA with multiple comparison correction test were performed with *, P<0.05; **, P<0.01. (**B**) mNG-positive cells at 24 and 48 h post-infection from (**A**). Scale bar, 100 μm. (**C**) RT-PCR analysis for single-round infection. For confirming the P10 ΔORF3-E virion remains infectious for only a single round on Vero cells, Vero E6 or Vero-ORF3-E cells were infected with WT mNG SARS-CoV-2 or P10 ΔORF3-E mNG virion for two rounds as described in **Fig. 1G**. Viral RNAs were extracted from the second-round culture fluids and analyzed by RT-PCR. The RT-PCR product, fragment 8 (gF8), is indicated in **Fig. 1B**. (**D**) Adaptive mutations. Three mutations were identified from whole genome sequencing of P10 ΔORF3-E mNG virion. No mutation was found in the P1 ΔORF3-E mNG virion.

### Exclusion of WT SARS-CoV-2 production

To confirm that no WT SARS-CoV-2 is inadvertently produced during *trans*-complementation, we performed four additional selections by passaging ΔORF3-E mNG virions on Vero-ORF3-E cells for five rounds. The P5 virions from selections I-III could only infect Vero cells for single round (**Fig. S4A-B**). Unexpectedly, selection IV produced P5 (S-IV-P5) virions that could infect parental Vero E6 cells for more than one round, though at a barely detectable level of ~10^2^ TCID_50_/ml, which was >100,000-fold lower than the WT mNG SARS-CoV-2 titers (**Fig. S4C**). To remove the single-round virion from the multi-round virion in the S-IV-P5 stock, we passaged the S-IV-P5 virion stock on Vero E6 cells for two rounds, resulting in S-IV-P5-Vero-P2 virion capable of multi-round infection. Full-genome sequencing revealed that the S-IV-P5-Vero-P2 virion retained the ORF3-E deletion but accumulated mutations in nsp15, nsp16, S, and M genes (**Fig. S4D**). Engineering the accumulated mutations into ΔORF3-E mNG RNA showed that the M mutation T130N conferred multi-round infection on Vero cells (**Fig. S4E**). Residue T130 is predicted to be on the intra-virion side of the M protein (*16, 17*), and is conserved in SARS-CoV and SARS-CoV-2 (**Fig. S4F**). The results indicate that, despite an absence of WT SARS-CoV-2 production and a lack of ORF and E genes, the *trans*-complementation system could produce mutant virions capable of infecting parental Vero cells for multiple rounds at a barely detectable level.

Next, we continuously cultured the S-IV-P5 variant on parental Vero E6 cells for 10 rounds (3-4 days per round) to select for potential virions with improved replication efficiency. However, passage did not improve viral replication on Vero cells (**Fig. S5**). The result suggests that, due to the lack of ORF3 and E gene, the S-IV-P5 virion is unlikely to gain efficient multiple-round amplification on normal cells through adaptation.

### Safety evaluation of ΔORF3-E virions *in vivo*

We examined the virulence of ΔORF3-E mNG virion in hamsters and K18-hACE2 transgenic mice (*18-20*). After intranasal inoculation with 6×10^5^ TCID_50_ of ΔORF3-E mNG virion (the highest possible infecting dose; **Fig. 3A**), hamsters did not lose weight (**Fig. 3B**) or develop detectable disease (**Fig. 3C**). In contrast, 10^5^ TCID_50_ of WT SARS-CoV-2-infected hamsters developed weight loss and mild disease (*e*.*g*., ruffled fur). The ΔORF3-E mNG virion-infected hamsters contained low levels of viral RNA in nasal washes (**Fig. 3D**) and oral swabs (**Fig. 3E**). Viral RNA levels in the trachea and lungs from the ΔORF3-E virion-infected animals were 5,000- and 400-fold lower than those from the WT virus-infected hamsters, respectively (**Fig. 3F**). Next, we examined the S-IV-P5-Vero-P2 virion, capable of infecting Vero cells for multiple rounds, in hamsters. To maximize the infection dose of S-IV-P5-Vero-P2 virion, we amplified S-IV-P5-Vero-P2 on Vero-ORF3-E cells, producing a virion stock of 5×10^4^ TCID_50_/ml. After intranasal inoculation with 5×10^3^ TCID_50_ of S-IV-P5-Vero-P2 virion (the highest possible dose), hamsters did not lose weight or develop detectable disease (**Fig. S6**). Collectively, the results indicate that both ΔORF3-E mNG virion and S-IV-P5-Vero-P2 virion are highly attenuated and do not disseminate or cause disease in hamsters.

**Figure 3.**
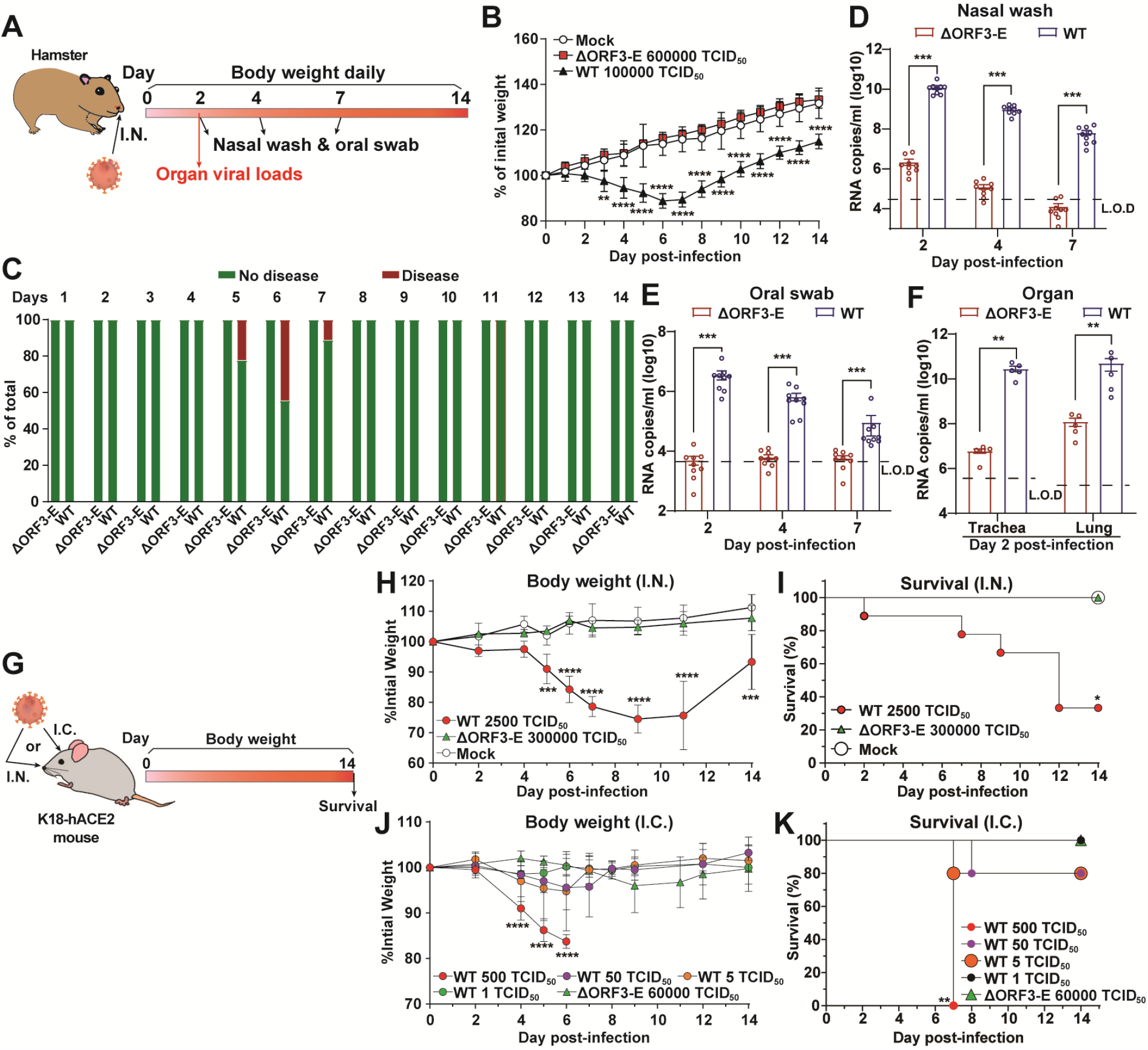
Safety characterization of ΔORF3-E mNG virion in animal models. (**A**) Hamster experimental schedule. Four- to five-week-old male Syrian golden hamsters were intranasally (I.N.) inoculated with 10^5^ TCID_50_ of WT SARS-CoV-2, 6×10^5^ TCID_50_ of ΔORF3-E mNG virion, or PBS mock control. Hamsters were monitored for weight loss, disease, and viral RNA level. (**B**) Hamster weight change (n=9). (**C**) Hamster disease (n=9). (**D**) Hamster nasal wash viral RNA level (n=9). (**E**) Hamster oral swab viral RNA level (n=9). (**F**) Viral RNA loads in hamster trachea and lung at day 2 post-infection (n=5). Limit of detection (L.O.D.) was defined as the RNA copies detected from mock-infected hamster samples. The weight loss data are shown as mean ± standard deviation and statistically analyzed using two-way ANOVA Turkey’s multiple comparison. The genomic RNA levels are presented as mean±standard error of the mean and analyzed by Mann-Whitney test. *, P<0.05; **, P<0.01; ***, P<0.001; ****, P<0.0001. (**G**) Mouse experimental schedule. Seven- to nine-week-old K18-hACE2 mice were inoculated with WT SARS-CoV-2 or ΔORF3-E mNG virion via the intranasal (I.N.) or intracranial (I.C.) route. Mouse weight loss after I.N. **(H)** or I.C. **(J)** infection. Body weights were normalized to the initial weight. The means for each group [I.N: WT SARS-CoV-2 (n=9), ΔORF3-E mNG virion (n=4), and mock (n=4); I.C: WT SARS-CoV-2 500 TCID_50_ (n=4), 50 TCID_50_ (n=5), 5 TCID_50_ (n=5), and 1 TCID_50_(n=5) and 6×10^4^ TCID_50_ ΔORF3-E mNG virus (n=4)] are indicated, with error bars indicating the standard deviation. A mixed-model ANOVA using Dunnett’s test for multiple comparisons was used to evaluate the statistical significance among groups: *, P<0.05; **, P<0.01; ***, P<0.001; ****, P<0.0001. Mouse survival after I.N. (**I**) and I.C. **(K)** inoculation was analyzed using the Gehan-Breslow-Wilcoxon test. Using groups with 100% survival as a comparator, a Bonferroni correction was applied manually to adjust the threshold for significance (indicated by *).

To corroborate the hamster results, we tested ΔORF3-E mNG virion in more susceptible K18-hACE2 mice (**Fig. 3G**). After intranasal inoculation with 3×10^5^ TCID_50_ of ΔORF3-E mNG virion (the highest possible dose), K18-hACE2 mice did not lose weight (**Fig. 3H**) or die (**Fig. 3I**); in contrast, infection with 2.5×10^3^ TCID_50_ of WT SARS-CoV-2 resulted in 25% weight loss and 67% lethality. To increase the stringency of the test, we inoculated K18-hACE2 mice by intracranial injection with 6×10^4^ TCID_50_ of ΔORF3-E mNG virion (the highest possible dose); no morbidity (**Fig. 3J**) or mortality (**Fig. 3K**) was observed. In contrast, mice inoculated by the intracranial route with 500, 50, 5, and 1 TCID_50_ of WT SARS-CoV-2 developed 100%, 25%, 25%, and 0% mortality, respectively (**Fig. 3K**). Similar to the ΔORF3-E mNG virion, no morbidity or mortality was observed after mice were inoculated by the intranasal or intracranial route with 2.5×10^3^ or 5×10^2^ TCID_50_ of S-IV-P5-Vero-P2 virion, respectively (**Fig. S7**). Together, the results demonstrate that both single-round ΔORF3-E mNG virion and multiple-round S-IV-P5-Vero-P2 virion lack virulence in K18-hACE2 mice.

### High-throughput neutralization and antiviral testing

We adapted ΔORF3-E mNG virion for a high-throughput neutralization and antiviral assay. **Fig. 4A** outlines the assay scheme in a 96-well plate format. Neutralization titers of 18 convalescent sera from COVID-19 patients were measured by two assays for comparison: the ΔORF3-E mNG virion assay and the gold standard plaque-reduction neutralization test (PRNT). The two assays produced comparable 50% neutralization titers (NT_50_) for all specimens (**Table S1 and Fig. 4B-C**). In addition, the ΔORF3-E mNG virion assay could also be used to measure the 50% effective neutralizing concentration (EC_50_) for a monoclonal antibody against SARS-CoV-2 receptor-binding domain (RBD; **Fig. 4D**). Finally, using Remdesivir as a viral polymerase inhibitor, we evaluated the ΔORF3-E mNG virion assay for antiviral testing. Remdesivir exhibited a more potent EC_50_ on hACE2-A549 cells (0.27 µM; **Fig. 4E**) than that on Vero cells (5.1 µM; **Fig. 4F**). The EC_50_ discrepancy between the two cell types is likely due to different efficiencies in converting Remdesivir to its triphosphate form, as previously reported (*3, 21*). Collectively, the results demonstrate that the ΔORF3-E virion assay can be used for high-throughput neutralization testing and antiviral drug discovery.

**Figure 4.**
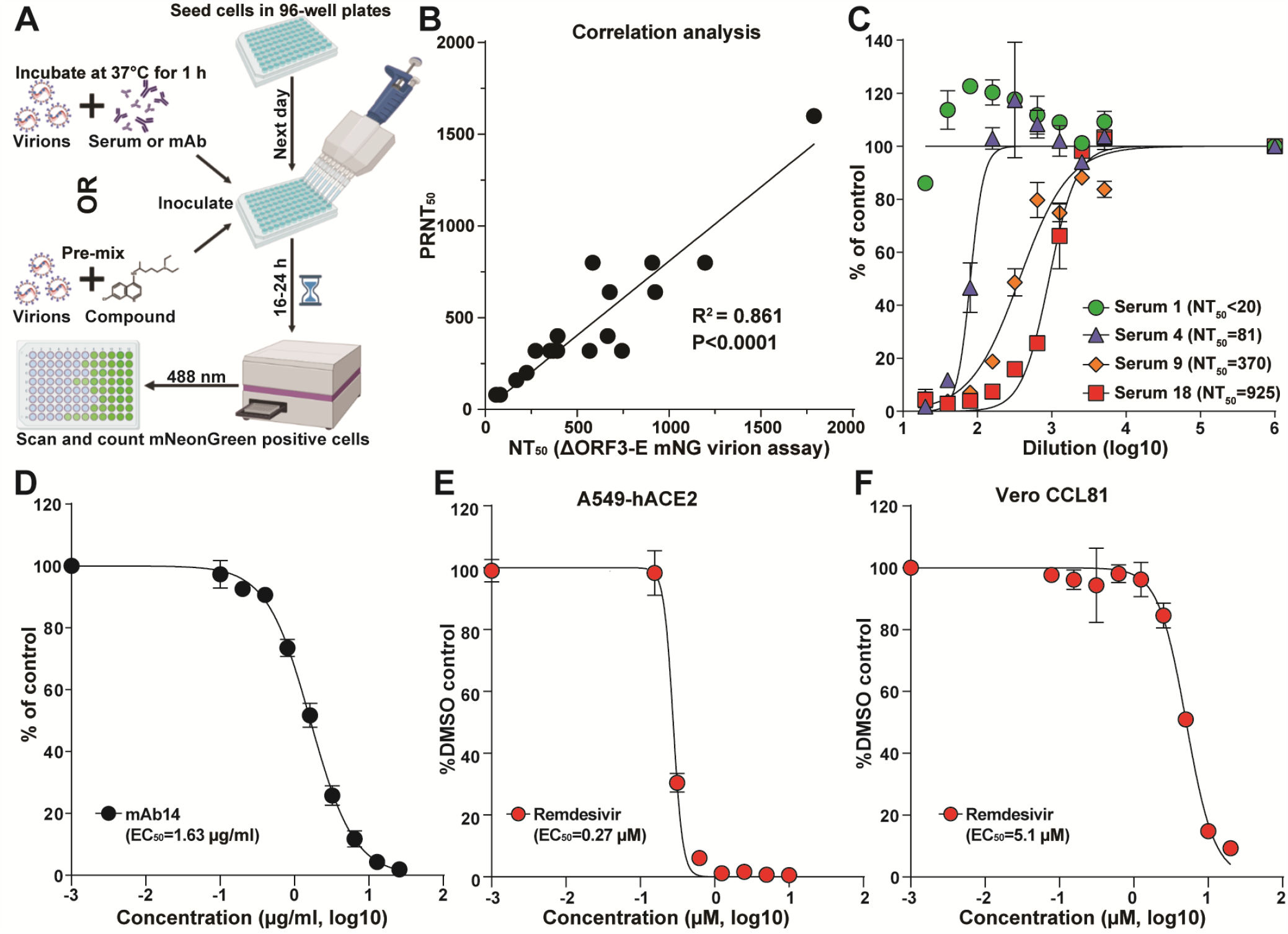
ΔORF3-E mNG virion-based high-throughput neutralization and antiviral testing. (**A**) Assay scheme in a 96-well format. (**B**) Correlation analysis of NT_50_ values between the ΔORF3-E mNG virion assay and plaque-reduction neutralization test (PRNT). The Pearson correlation efficiency R^2^ and P values (two-tailed) are indicated. (**C**) Neutralization curves. Representative curves are presented for one negative and three positive sera. The means and standard deviations from two independent experiments are shown. (**D**) EC_50_ of human mAb14 against ΔORF3-E mNG virion infecting Vero CCL81 cells. The mean ± standard deviations from four independent experiments are indicated. (**E**) EC_50_ of Remdesivir against ΔORF3-E mNG virion infecting A549-hACE2 cells. (**F**) EC_50_ of Remdesivir against ΔORF3-E mNG virion on Vero CCL81 cells. For (**E**) and (**F**), the mean ± standard deviations from three independent experiments are indicated. The four-parameter dose-response curve was fitted using the nonlinear regression method.

## DISCUSSION

We generated and characterized a *trans*-complementation system for SARS-CoV-2. The system produced a high yield of single-round infectious ΔORF3-E virion that could be used for neutralization and antiviral testing. An mNG reporter was introduced into the ΔORF3-E virion to indicate viral replication. Depending on research needs, other reporter genes, such as luciferase or GFP, could be engineered into the system. A reliable high-throughput neutralization assay is important for COVID-19 vaccine evaluation and for studying the kinetics of neutralizing antibody levels in post-vaccinated and naturally infected people (*4, 22, 23*). Three types of cell-based high-throughput neutralization assays currently are available: (i) pseudovirus assay, which expresses SARS-CoV-2 S protein alone on heterologous viruses, can be performed at BSL-2 laboratories (*24, 25*); (ii) a reporter SARS-CoV-2 assay, which must be performed at BSL-3 laboratories, represents authentic viral infection (*3, 5, 6, 26*); (iii) *bona fide* fully infectious SARS-CoV-2 by focus reduction neutralization test (*24*). The ΔORF3-E mNG virion combines the advantages of each assay type by recapitulating the authentic viral infection for a single round, thus supporting its use at BSL2 laboratories. The ΔORF3-E mNG virion can be readily adapted to investigate vaccine-elicited neutralization against newly emerged SARS-CoV-2 isolates, such as the rapidly spreading United Kingdom and South African strains (*27, 28*), by swapping or mutating the S gene.

The *trans-*complementation system can also be used for high-throughput antiviral screening of large compound libraries. Infection of normal cells with ΔORF3-E mNG virion allows for screening of inhibitors of virus entry, genome translation, and RNA replication, but not virion assembly/release. In contrast, infection of Vero-ORF3-E cells with ΔORF3-E mNG virion can be used to identify inhibitors of all steps of SARS-CoV-2 infection cycle, including virion assembly and release; this system also allows selection for resistance to inhibitors for mode-of-action studies. In addition, the single-round ΔORF3-E virion could be developed as a safe vaccine platform, as previously reported for other coronaviruses (*29, 30*).

Our results support that the *trans*-complementation system can be performed safely in BSL-2 laboratories. (i) The system produced single-round infectious ΔORF3-E mNG virion that does not infect normal cells for multiple rounds and thus cannot spread *in vitro* or *in vivo*. (ii) The system did not produce WT virus, even after multiple independent selections. (iii) Although an adaptive mutation in M protein was selected to confer multi-round infection on normal cells, the replication level of such virion (*i*.*e*., S-IV-P5-Vero-P2) was barely detectable, with infectious titers >100,000-fold lower than the WT SARS-CoV-2. The molecular mechanism of how S-IV-P5-Vero-P2 could infect cells for multiple rounds without the ORF3 and E proteins remains to be defined. Previous studies showed that deletion of both ORF3 and E genes was lethal for SARS-CoV (*31*). (iv) Continuous culturing of the S-IV-P5-Vero-P2 virion on naïve Vero cells did not improve viral replication. (v) When hamsters and K18-hACE2 mice were infected with the highest possible doses, neither ΔORF3-E mNG virion nor S-IV-P5-Vero-P2 virion caused morbidity or mortality. Even after intracranial infection with the highest possible dose, neither virions caused detectable disease or death in the highly susceptible K18-hACE2 mice. If further safety improvement is needed, more accessory ORFs could be deleted from the ΔORF3-E mNG RNA as accessory proteins are not essential for viral replication (*1*).

One limitation of our study is the use of Vero E6 cells for constructing the Vero-ORF3-E cell line. When propagated on Vero E6 cells, SARS-CoV-2 could accumulate deletions at the furin cleavage site in the S protein (*32, 33*). This cleavage deletion affects the neutralization susceptibility of SARS-CoV-2 and possibly the route of entry into cells (*34*). Although we did not observe furin cleavage deletions when our ΔORF3-E mNG virion was passaged on Vero-ORF3-E cells, this possibility could be minimized or eliminated by using other cell lines, such as A549-hACE2 or Vero-TMPRSS2-hACE2 cells.

In summary, we have developed a *trans*-complementation system for SARS-CoV-2 that likely can be performed at BSL-2 laboratories for COVID-19 research and countermeasure development. Thus, the experimental system could be used by researchers in industry, academia, and government laboratories who lack access to a BSL-3 facility.

## METHODS

### Cell lines

Vero E6, Vero CCL-81, Calu-3, and HEK-293T cells were purchased from the American Type Culture Collection (ATCC) and cultured in high-glucose Dulbecco’s modified Eagle’s medium (DMEM) supplemented with 2 mM L-glutamine, 100 U/ml Penicillium-Streptomycin (P/S), and 10% fetal bovine serum (FBS; HyClone Laboratories, South Logan, UT). Vero-ORF3-E cells were maintained in DMEM medium supplemented with 2mM L-glutamine, 100 U/ml P/S, 10% FBS, 0.075% sodium bicarbonate, and 10 μg/ml puromycin. The A549-hACE2 cells were generously provided by Shinji Makino (*35*) and grown in the culture medium supplemented with 10 μg/mL blasticidin and 10 mM HEPES at 37 °C with 5% CO2. Medium and other supplements were purchased from Thermo Fisher Scientific (Waltham, MA).

### Hamsters

Syrian golden hamsters (HsdHan:AURA strain) were purchased from Envigo. Animals were housed in groups and fed standard chow diets. Hamster experiments were performed as described previously (*36*). Briefly, 10^5^ TCID_50_ of WT SARS-CoV-2, 6×10^5^ TCID_50_ of ΔORF3-E mNG virion, or 5×10^3^ TCID_50_ of S-IV-P5-Vero-P2 virion in 100 μl volume were inoculated into four- to five-week-old male Syrian golden hamsters via the intranasal route. The S-IV-P5-Vero-P2 virion stock was prepared by two rounds of culturing of S-IV-P5 virion (**Fig. S4A**) on Vero E6 cells (to remove single-round infectious virion), followed by propagation on Vero-ORF3-E cells. Fourteen hamsters were used in SARS-CoV-2- and ΔORF3-E mNG virion-infected groups and 5 hamsters were used in S-IV-P5-Vero-P2 virion-infected group. From day 1 to 14 post-infection, hamsters were observed daily for weight change and signs of illness. Five hamsters in WT SARS-CoV-2-, ΔORF3-E mNG virions-, or mock-infected group were sacrificed on day 2 post-infection for lung and trachea collections. Nasal washes and oral swabs of the rest 9 hamsters per group were collected on days 2, 4, and 7 post-infection.

### Mice

Animal studies were carried out in accordance with the recommendations in the Guide for the Care and Use of Laboratory Animals of the National Institutes of Health. The protocols were approved by the Institutional Animal Care and Use Committee at the Washington University School of Medicine (assurance number A3381–01). Heterozygous K18-hACE c57BL/6J mice (strain: 2B6.Cg-Tg(K18-ACE2)2Prlmn/J) were obtained from the Jackson Laboratory. Animals were randomized upon arrival at Washington University and housed in groups of <5 per cage in rooms maintained between 68-74°F with 30-60% humidity and day/night cycles of 12 h intervals (on 6AM-6PM). Mice were fed standard chow diets. Mice 7-9 weeks of age and of both sexes were used for this study. Intranasal virus inoculations (50 uL/mouse) were performed under sedation with ketamine hydrochloride and xylazine while intracranial virus inoculations (10 μL/mouse) were performed under sedation with isoflurane; all efforts were made to minimize animal suffering.

### Plasmid construction

Seven previously reported subclone plasmids for the assembly of the entire genome of SARS-CoV-2 were used in this study, including pUC57-F1, pCC1-F2, pCC1-F3, pUC57-F4, pUC57-F5, pUC57-F6, and pCC1-F7-mNG (*2, 26*). For the convenience of deleting ORF3-E gene, we constructed F5, F6, and F7-mNG fragments into one plasmid. F5, F6, and F7-mNG fragments were amplified from corresponding subclones via PCR with primer pairs pcov-F56-F1/pncov-R5, pncov-F6/pncov-R6, and pncov-F7/pncov-R8, respectively (**Table S2**). All PCR products were cloned together into a pCC1 vector through NotI and ClaI restriction sites using the standard restriction digestion-ligation cloning, resulting in subclone pCC1-F567-mNG.

To introduce ORF3-E deletion and mutant Transcription Regulatory Sequence (TRS) into pCC1-F567-mNG, seven fragments were amplified with primer pairs cov-21115-F/TRS2-S-R, TRS2-S-F/S-TRS2-M-R, TRS2-M-F/M-TRS2-R, M-TRS2-F/ORF6-TRS2-mNG-R, ORF6-TRS2-mNG-F/ORF7-TRS2-ORF8-R, ORF7-TRS2-ORF8-F/ORF8-TRS2-N-R, and TRS2-N-F/cov-28501-R. The seven PCR products were assembled into the pCC1-F567-mNG plasmid that were pre-linearized with NheI and XhoI by using the NEBuilder® HiFi DNA Assembly kit (NEB) according to the manufacturer’s instruction, resulting in subclone pCC1-F567-mNG-ΔORF3-E. Mutation T130N in M protein was engineered into pCC1-F567-mNG-ΔORF3-E with primers M-T130N-F/M-T130N-R via overlap PCR. Mutant TRS was engineered into pCC1-F1 with primers 5□UTR-TRS2-F and 5□UTR-TRS2-R via overlap PCR.

For making the Vero-ORF3-E cell lines, codon-optimized SARS-CoV-2 ORF3 and E genes were synthesized by GenScript Biotech (Piscataway, NJ). An mCherry reporter Zika virus cDNA plasmid (*37*) was used as a template to amplify the mCherry-F2A gene. For constructing a lentiviral plasmid expressing ORF3 and E protein of SARS-CoV-2, DNA fragments encoding mCherry-F2A, SARS-CoV-2 E, EMCV IRES, and SARS-CoV-2 ORF3 were amplified with primers EcoR1-mCherry-F/F2A-optE-R, F2A-optE-F/EcoR1-Cov-optE-R, EcoR1-IRES-F/EMCV-IRES-R, and IRES-optORF3-F/BamH1-Cov-optORF3-R, respectively. The PCR products then were inserted into a Tet-on inducible lentiviral vector pLVX (Takara, Mountain View, CA) through EcoRI and BamHI restriction sites, resulting in plasmid pLVX-ORF3-E.

### Selection of Vero-ORF3-E cell line

For packaging the lentivirus, the pLVX-ORF3-E plasmid was transfected into HEK-293T cells using the Lenti-X Packaging Single Shots kit (Takara). Lentiviral supernatants were harvested at 72 h post-transfection and filtered through a 0.22 μM membrane (Millipore, Burlington, MA). One day before transduction, Vero E6 cells were seeded in a 6-well plate (4×10^5^ per well) with DMEM containing 10% FBS. After 12-18 h, cells were transduced with 2 ml lentivirus for 24 h in the presence of 12 μg/ml of polybrene (Sigma-Aldrich, St. Louis, MO). At 24 h post-transduction, cells from a single well were split into four 10 cm dishes and cultured in medium supplemented with 25 μg/ml of puromycin. The culture medium containing puromycin was refreshed every 2 days. After 2-3 weeks of selection, visible puromycin-resistant cell colonies were formed. Several colonies were transferred into 24-well plates. When confluent, cells were treated with trypsin and seeded in 6-well plates for further expansion. The resulting cells were defined as Vero-ORF3-E P0 cells. For cell line verification, total cellular mRNA was isolated and subject to RT-PCR with primers EcoR1-mCherry-F and BamH1-Cov-optORF3-R (**Table S2**), followed by cDNA sequencing of the ORF3-E genes.

### ΔORF3-E mNG cDNA assembly and *in vitro* RNA transcription

Full-length genome assembly and RNA transcription were performed as described previously with minor modifications (*2*). Briefly, individual subclones containing fragments of the ΔORF3-E mNG viral genome were digested with appropriated restriction endonucleases and resolved in a 0.8% agarose gel. Specifically, the plasmids containing F1, F2, F3, or F4 fragments were digested with BsaI enzyme, and the plasmid containing F567-mNG-ΔORF3-E fragment was digested with EspI enzyme. All fragments were recovered using the QIAquick Gel Extraction Kit (QIAGEN, Hilden, Germany), and total of 5 μg of the five fragments was ligated in an equal molar ratio by T4 DNA ligase (New England Biolabs, Ipswich, MA) at 4°C overnight. Afterward, the assembled full-length genomic cDNA was purified by phenol-chloroform extraction and isopropanol precipitation. ΔORF3-E mNG RNA transcripts were generated using the T7 mMessage mMachine kit (Ambion, Austin, TX). To synthesize the N gene RNA transcript of SARS-CoV-2, the N gene was PCR-amplified by primers CoV-T7-N-F and polyT-N-R (**Table S2**) from a plasmid containing the F7 fragment (*2*); the PCR product was then used for *in vitro* transcription using the T7 mMessage mMachine kit (Ambion).

### ΔORF3-E mNG virion production and quantification

Vero-ORF3-E cells were seeded in a T175 flask and grown in DMEM medium with 100 ng/ml of doxycycline. On the next day, 40 μg of ΔORF3-E mNG RNA and 20 μg of N-gene RNA were electroporated into 8×10^6^ Vero-ORF3-E cells using the Gene Pulser XCell electroporation system (Bio-Rad, Hercules, CA) at a setting of 270V and 950 μF with a single pulse. The electroporated cells were then seeded in a T75 flask and cultured in the medium supplemented with doxycycline (Sigma-Aldrich) at 37°C for 3-4 days. Virion infectivity was quantified by measuring the TCID_50_ using an end-point dilution assay as previously reported (*38*). Briefly, Vero-ORF3-E cells were plated on 96-well plates (1.5×10^4^ per well) one day prior to infection. The cells were cultured in medium with doxycycline as described above. ΔORF3-E mNG virions were serially diluted in DMEM medium supplemented with 2% FBS, with 6 replicates per concentration. Cells were infected with 100 μl of diluted virions and incubated at 37°C for 2-3 days. The mNG signals were counted under a fluorescence microscope (Nikon, Tokyo, Japan). TCID_50_ was calculated using the Reed & Muench method (*39*).

To assess viral RNA levels, a quantitative RT-PCR assay was conducted using an iTaq Universal SYBR Green one-step kit (Bio-Rad) on a QuantStudio 7 Flex Real-Time PCR Systems (Thermo fisher) by following the manufacturers’ protocols. Primers CoV19-N2-F and CoV19-N2-R (**Table S2**) targeting the N gene were used. Absolute RNA copies were determined by standard curve method using *in vitro* transcribed RNA containing genomic nucleotide positions 26,044 to 29,883 of the SARS-CoV-2 genome.

### RNA extraction, RT-PCR, and cDNA sequencing

Supernatants of infected cells were collected and centrifuged at 1,000 g for 10 min to remove cell debris. Clarified culture fluids (250 μl) were mixed thoroughly with 1 ml of TRIzol LS reagent (Thermo Fisher Scientific). Extracellular RNA was extracted per manufacture’s instruction and resuspended in 20 μl of nuclease-free water. RT-PCR was performed using the SuperScript® IV One-Step RT-PCR kit (Thermo Fisher Scientific). Nine cDNA fragments (gF1 to gF9) covering the whole viral genome were generated with specific primers according to the protocol described previously (*2*). Afterward, cDNA fragments were separated in a 0.8% agarose gel, purified using QIAquick Gel Extraction Kit (QIAGEN), and subjected to Sanger sequencing.

### ΔORF3-E mNG virion neutralization assay

The research protocol for use of human serum specimens was approved by the University of Texas Medical Branch (UTMB) Institutional Review Board (IRB protocol number 20-0070). All human serum specimens were obtained at the UTMB with patient information de-identified. For neutralization testing, Vero CCL-81 cells (1.2×10^4^) in 50 μl of DMEM containing 2% FBS and 100 U/ml P/S were seeded in each well of black μCLEAR flat-bottom 96-well plate (Greiner Bio-one™, Kremsmünster, Austria). At 16 h post-seeding, 30 μL of 2-fold serial diluted human sera were mixed with 30 μL of ΔORF3-E mNG virion (MOI of 5) and incubated at 37°C for 1 h. Afterward, 50 μL of virus–sera complexes were transferred to each well of the 96-well plate. After incubating the infected cells at 37°C for 20 h, 25 μl of Hoechst 33342 Solution (400-fold diluted in Hank’s Balanced Salt Solution; Thermo Fisher Scientific) were added to each well to stain the cell nucleus. The plate was sealed with Breath-Easy sealing membrane (Diversified Biotech, Dedham, MA), incubated at 37°C for 20 min, and quantified for mNG-positive cells using the CellInsight CX5 High-Content Screening Platform (Thermo Fisher Scientific). Infection rates were determined by dividing the mNG-positive cell number to the total cell number. Relative infection rates were obtained by normalizing the infection rates of serum-treated groups to those of non-serum-treated controls. The curves of the relative infection rates versus the serum dilutions (log10 values) were plotted using Prism 9 (GraphPad, San Diego, CA). A nonlinear regression method was used to determine the dilution fold that neutralized 50% of mNG fluorescence (NT_50_). Each serum was tested in duplicates.

### ΔORF3-E mNG virion for mAb and antiviral testing

Vero CCL-81 cells (1.2×10^4^) or A549-hACE2 cells in 50 μl of culture medium containing 2% FBS were seeded in each well of black μCLEAR flat-bottom 96-well plate. At 16 h post-seeding, 2- or 3-fold serial diluted human mAb14 (*40*) or Remdesivir were mixed with ΔORF3-E mNG virion (MOI of 1). Fifty microliters of mixtures were transferred to each well of the 96-well plate. After incubating the infected cells at 37°C for 20 h, 25 μl of Hoechst 33342 Solution (400-fold diluted in Hank’s Balanced Salt Solution) were added to each well to stain the cell nucleus. The plate was sealed with Breath-Easy sealing membrane, incubated at 37°C for 20 min. mNG-positive cells were quantified and infection rates were calculated as described above. Relative infection rates were obtained by normalizing the infection rates of treated groups to those of non-treated controls. For Remdesivir, 0.1% of DMSO-treated groups were used as controls. A nonlinear regression method was used to determine the concentration that inhibited 50% of mNG fluorescence (EC_50_). Experiments were performed in triplicates or quadruplicates.

### Biosafety

All aspects of this study were approved by the Institutional Biosafety Committee of the University of Texas Medical Branch at Galveston before the initiation of this study. Experiments with SARS-CoV-2, *trans-*complementation, and ΔORF3-E mNG virion were performed in a BSL-3 laboratory by personnel equipped with powered air-purifying respirators.

### Transmission Electron Microscopy

Supernatants of infected cells were centrifuged for 10 min at 3,000 g to remove cellular debris. Nickel grids were incubated with clarified supernatants for 10 min followed by glutaraldehyde fixation and 2% uranyl acetate staining. Micrographs were taken using a JEM 1400 (JEOL USA Inc.). Multiple randomly selected fields were imaged.

### Bioinformatics analysis

Fluorescence images were processed using ImageJ (*41*). Virus sequences were download from the NCBI database and aligned using Geneious software. DNA gel images were analyzed using Image Lab software. Statistical graphs or charts were created using the GraphPad Prism 9 software. Figures were created and assembled using BioRender and Adobe illustration (San Jose, CA).

### Statistical analysis

A linear regression model in the software Prism 9 (GraphPad) was used to calculate the NT_50_ and EC_50_ values from the ΔORF3-E virion assay. Pearson correlation coefficient and two-tailed p-value are calculated using the default settings in the software Prism 9. An unpaired T-test (for two-groups comparison) and ANOVA test (for multi-group comparison) were used in statistical analysis (*, P<0.05, significant; **, P<0.01, very significant; ***, P<0.001, highly significant; ****, P<0.0001, extremely significant; ns, P>0.05, not significant).

## DATA AVAILABILITY

The results presented in the study are available upon request from the corresponding authors. The mNG reporter SARS-CoV-2 has been deposited to the World Reference Center for Emerging Viruses and Arboviruses (https://www.utmb.edu/wrceva) at UTMB for distribution.

## ACKNOWLEDGEMENTS

We thank John Bilello from Gilead for providing Remdesivir and Zhiqiang An from the University of Texas Health Science at Houston for providing mAb14. P.-Y.S. was supported by NIH grants AI142759, AI134907, AI145617, and UL1TR001439; awards from the Sealy & Smith Foundation, Kleberg Foundation, John S. Dunn Foundation, Amon G. Carter Foundation, Gilson Longenbaugh Foundation, and Summerfield Robert Foundation; and fund in sponsored research agreement from Q^2^ Solutions. M.S.D. was supported by R01 AI157155. V.D.M. was supported by NIH grants U19AI100625, R00AG049092, R24AI120942, and a STARs Award from the University of Texas System. S.C.W. was supported by NIH grant R24 AI120942. J.L. is supported by the postdoctoral fellowship from the McLaughlin Fellowship Endowment at UTMB. P.R. and X.X. were partially supported by the Sealy & Smith Foundation.

## AUTHOR CONTRIBUTIONS

X.Z., V.D.M., X.X., and P.-Y.S conceived the study. X.Z., Y.L., J.L., A.L.B., K.S.P., J.A.P., J.Z., H.X., N.B., P.R., and X.X. performed the experiments. X.Z., Y.L., A.L.B., P.A., P.R., V.D.M., M.S.D., S.W., X.X., and P.-Y.S. analyzed the results. P.R. prepared the serum specimens. X.Z., Y.L., A.L.B., V.D.M., M.S.D., S.W., X.X., and P.-Y.S wrote the manuscript.

## SUPPLEMENTAL FIGURE LEGENDS

**Figure S1.**
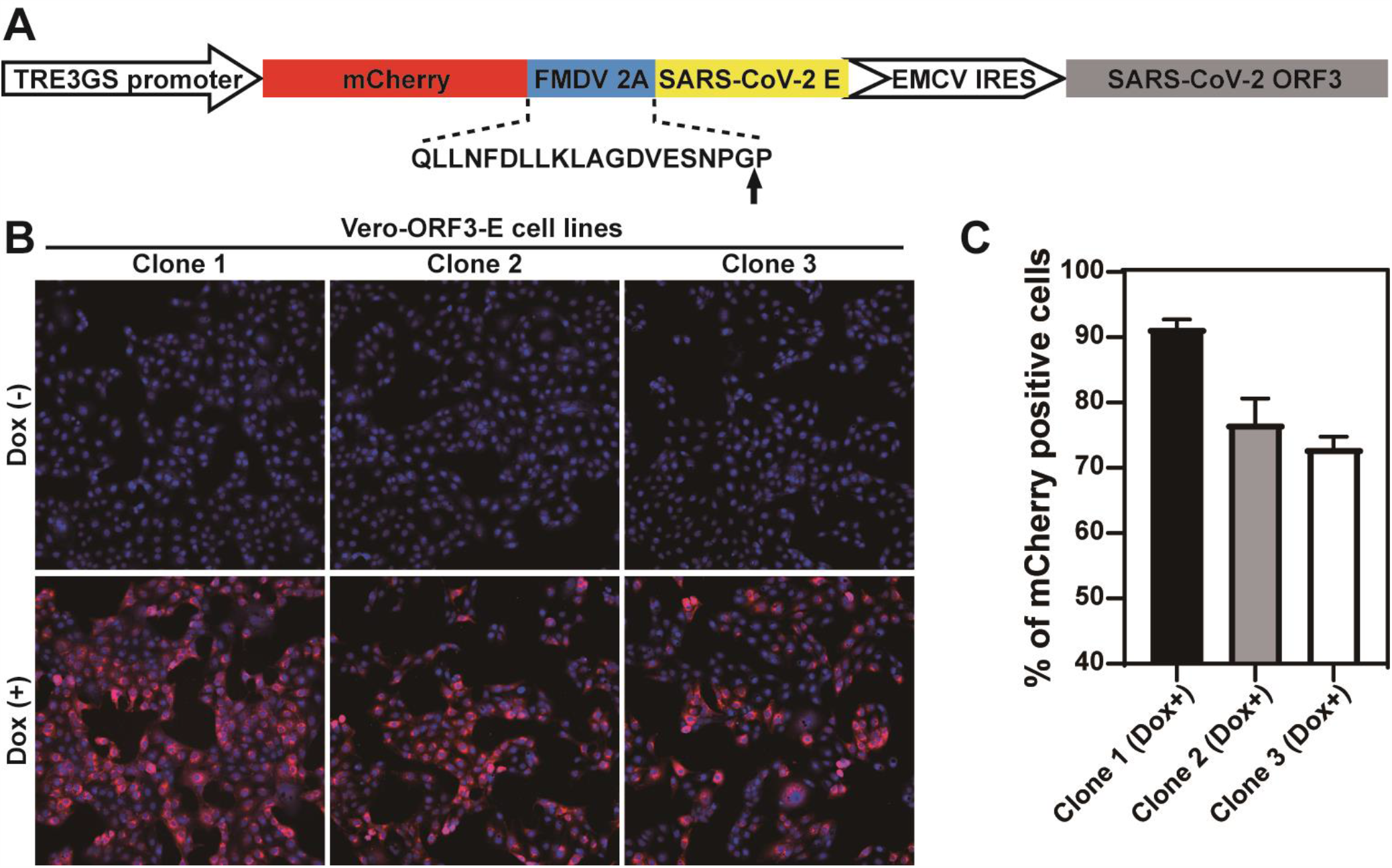
Construction of Vero-ORF3-E cell lines. (**A**) Construction of a lentiviral transfer plasmid encoding mCherry, ORF3, and E protein. The sequence of FMDV 2A and its translational break position is indicated by an arrow. (**B**) Merged mCherry (red) and nuclei (blue) images of 3 selected clones of Vero-ORF3-E cell lines. Nuclei were stained with Hoechst 33342. Doxycycline induction is indicated. (**C**) mCherry expression in doxycycline-induced cells. mCherry-positive cells were quantified using a plate reader. The percentages of mCherry positive cells are presented. The results are presented as means ± standard deviations from six replicates, and more than 10^5^ cells were counted for each clone. Clone 1 was used in the rest of this study.

**Figure S2.**
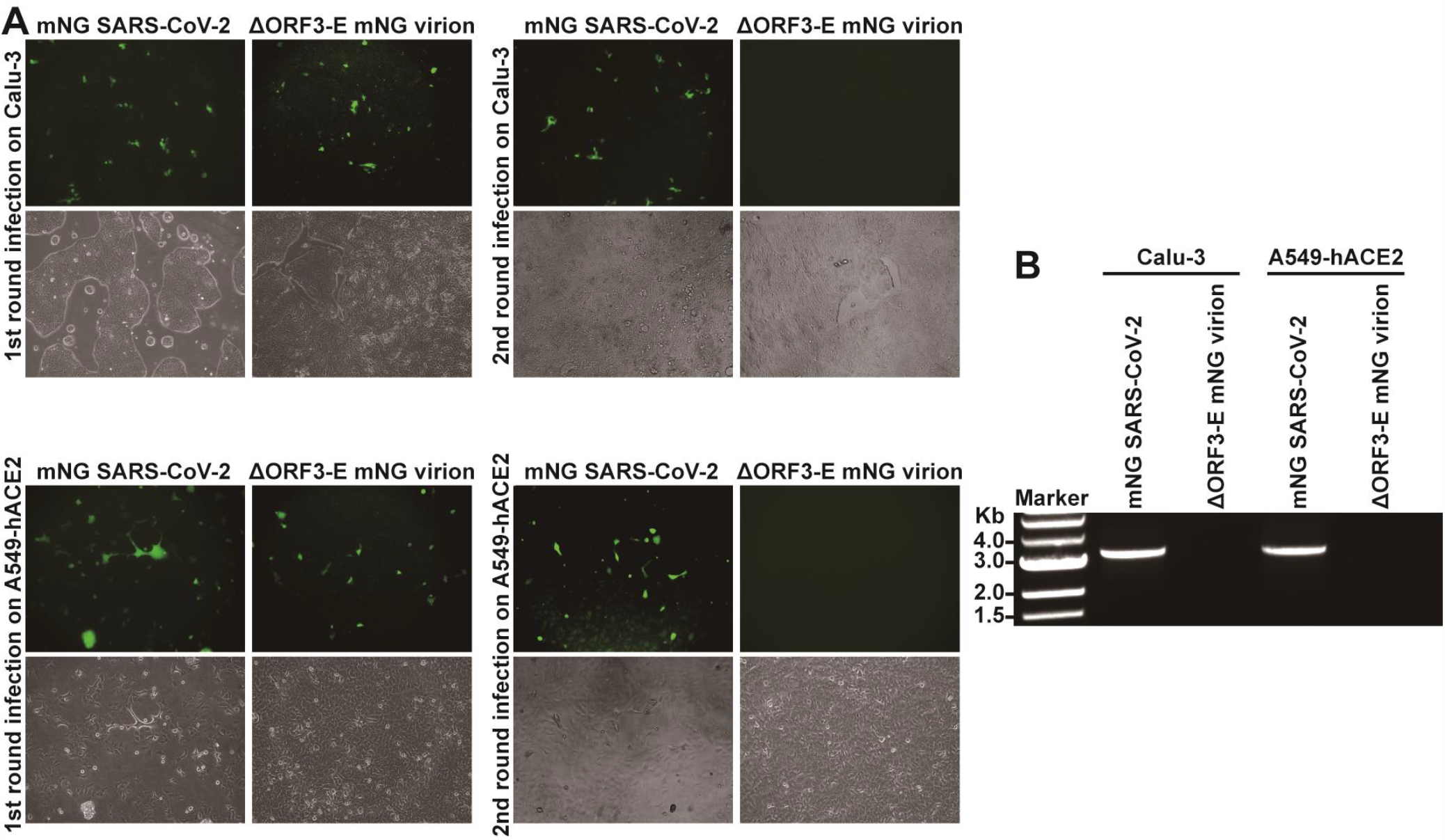
Single-round infection of ΔORF3-E mNG virion. (**A**) Calu-3 and A549-hACE2 cells (MOI of 1 and 10 for mNG SARS-CoV-2 and ΔORF3-E mNG virion, respectively; viral titers determined on Vero-ORF3-E cells) were infected with mNG SARS-CoV-2 or ΔORF3-E mNG virion for 2 h, after which the cells were washed and cultured in fresh medium. At day 2 post-infection, supernatants of the infected cells were transferred to infect naïve Calu-3 and A549-hACE2 for the second round. Fluorescence and phase contrast images for the infected cells are presented. (**B**) RT-PCR analysis of viral RNA. Extracellular RNAs from the second round of infection from (**A**) were harvested at day 2 post-infection and subjected to RT-PCR analysis of viral RNA.

**Figure S3.**
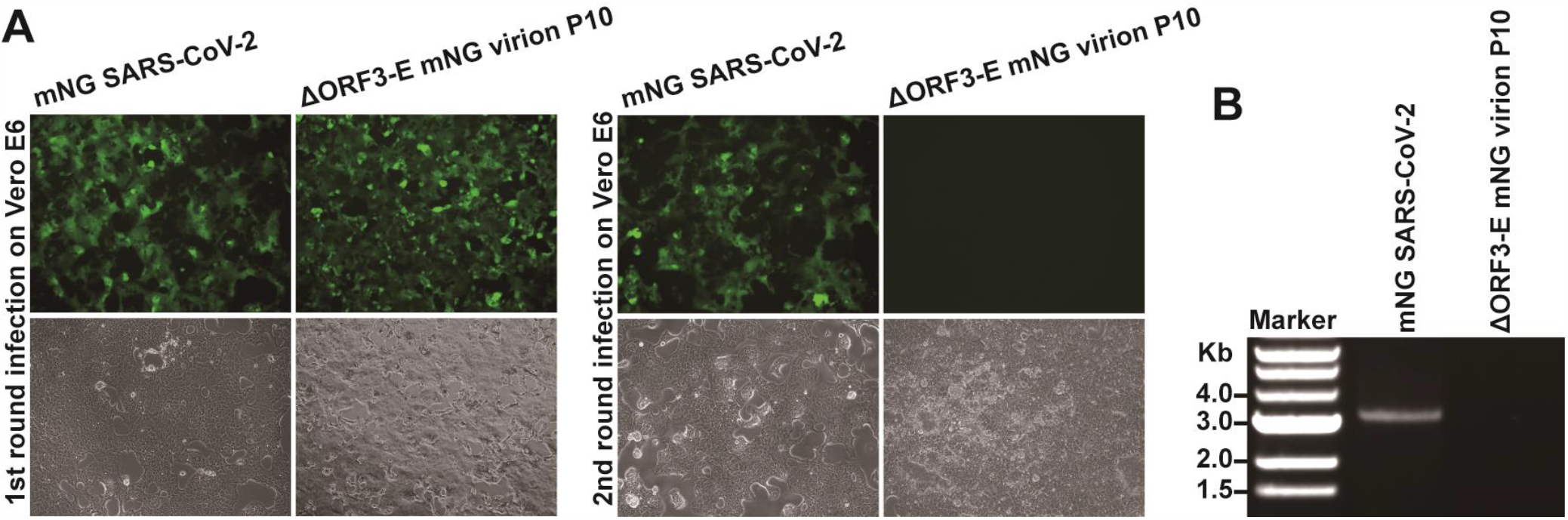
No WT mNG SARS-CoV-2 production from the *trans*-complementation system. WT mNG SARS-CoV-2 and P10 ΔORF3-E mNG virion (derived from 10 rounds of passaging of ΔORF3-E mNG virion on Vero-ORF3-E cells) were used to infect Vero E6 cells for two rounds as described in **Fig. 1G**. (**A**) Fluorescence and phase contrast images of infected cells are presented for both the first and second rounds of infections. (**B**) RT-PCR analysis of viral RNA extracted from the culture fluids from the second-round infected cells.

**Figure S4.**
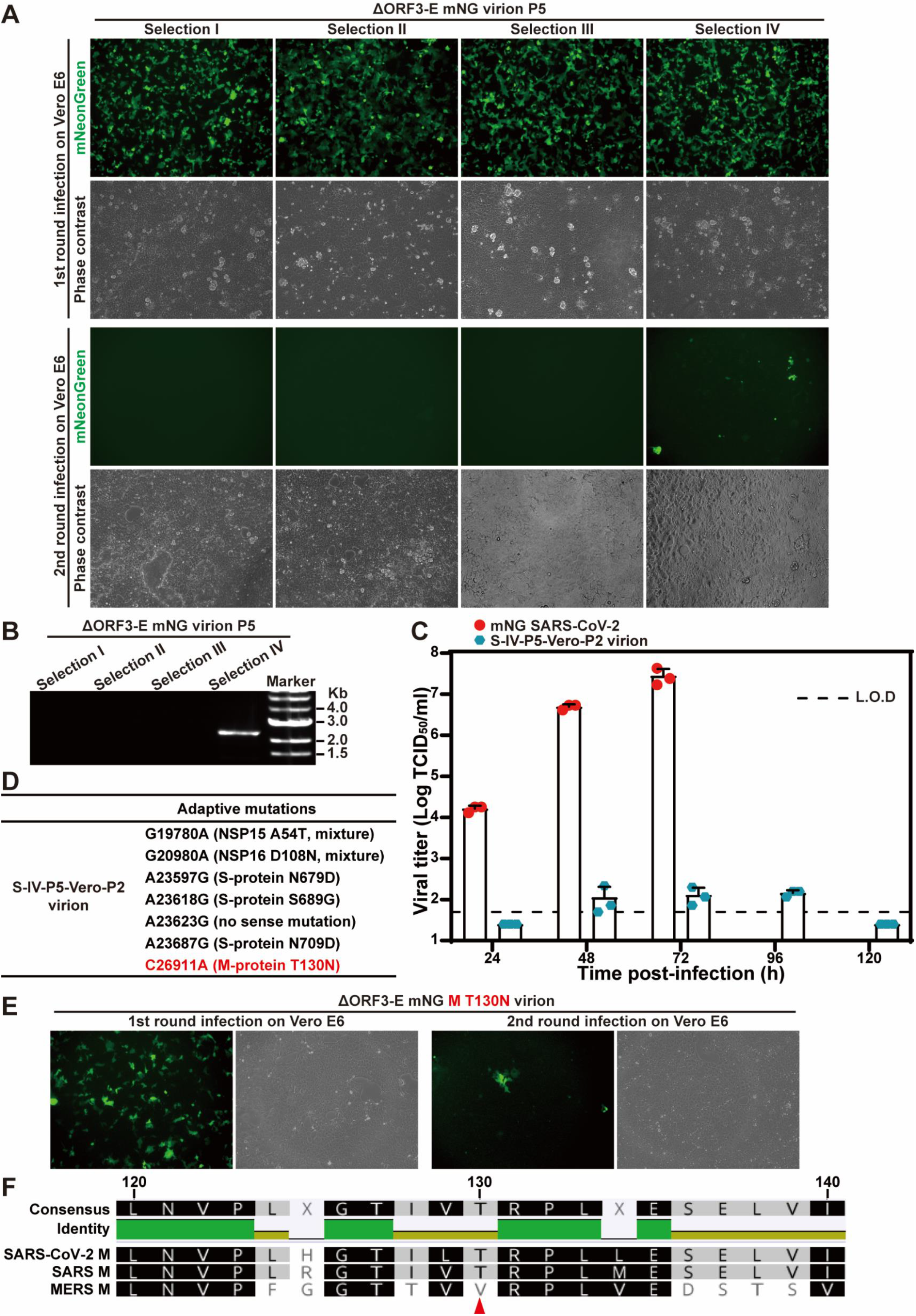
Selection of ΔORF3-E mNG virion capable of inefficiently infecting Vero E6 cells for more than one round. Four independently selected P5 ΔORF3-E mNG virions (generated from five rounds of passaging ΔORF3-E mNG virion on Vero-ORF3-E cells) were used to infect naïve Vero E6 cells for two rounds as described in **Fig. 1G**. The P5 ΔORF3-E mNG virion-infected Vero E6 cells were analyzed for mNG signals (**A**). The extracellular RNA from the second-round infected cells were examined for viral RNA by RT-PCR (**B**). Selection IV P5 ΔORF3-E (S-IV-P5) mNG virion could infect Vero cells for multiple rounds. To remove the single-round virion from the multi-round virion in the S-IV-P5 stock, the S-IV-P5 stock was passaged on Vero E6 cells for two rounds, resulting in S-IV-P5-Vero-P2 virion capable of multi-round infection. The replication kinetics of WT mNG SARS-CoV-2 and S-IV-P5-Vero-P2 mNG virion were compared on Vero E6 cells (**C**). The cells were inoculated at an MOI of 0.001. Limit of detection, L.O.D. Adaptive mutations were identified from the S-IV-P5-Vero-P2 mNG virion (**D**). The T130N mutation from the M protein was engineered into ΔORF3-E mNG virion. The resulting ΔORF3-E mNG M T130N virion was used to infect Vero E6 cells for two rounds. Fluorescence and phase contrast images of the infected cells are shown (**E**). Sequence alignment shows that the M proteins from SARS-CoV and SARS-CoV-2 share the same T130 residue (**F**). Red arrow indicates the T130 residue of SARS-CoV-2.

**Figure S5.**
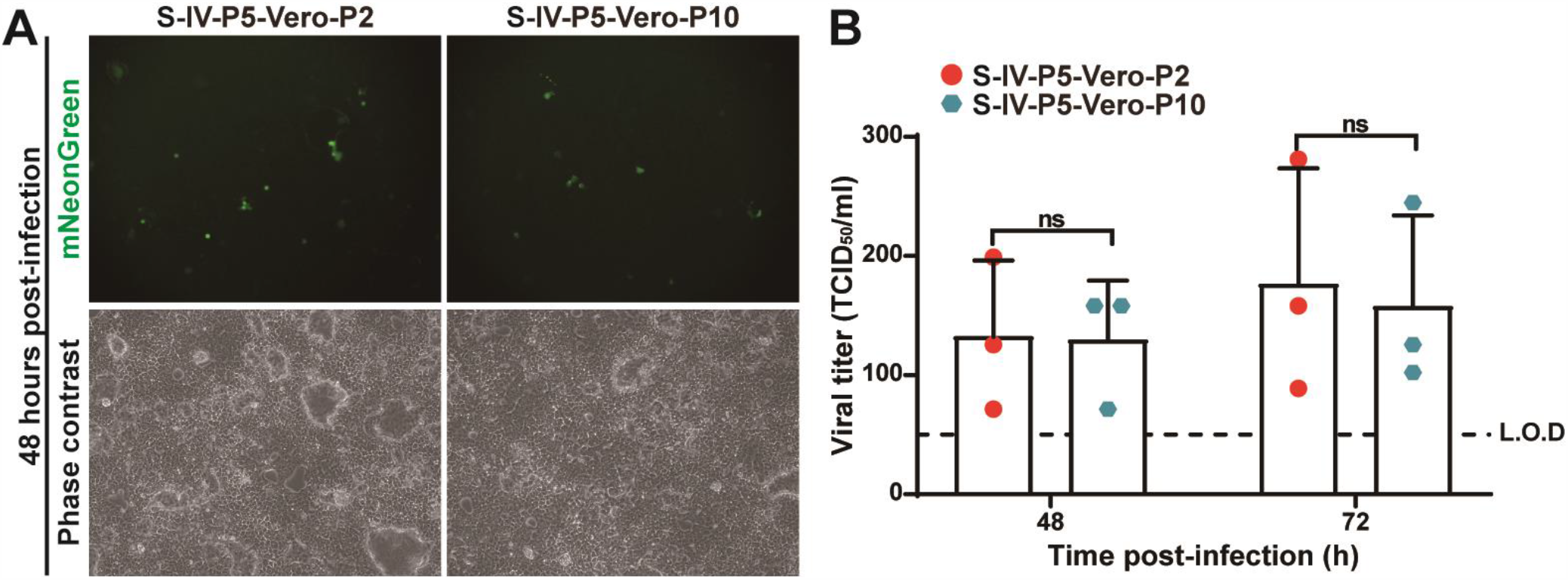
No improvement of viral replication of selection IV ΔORF3-E (S-IV-P5) mNG virion after 10 rounds of culturing on Vero E6 cells. S-IV-P5 mNG virion was continuously passaged on Vero E6 cells for 10 rounds. The resulting P2 and P10 S-IV-P5 mNG virions (*i*.*e*., S-IV-P5-Vero-P2 and S-IV-P5-Vero-P10, respectively) were used to infect Vero E6 cells at an MOI of 0.001. The mNG-positive cells (**A**) and the growth kinetics of the S-IV-P5-Vero-P2 and S-IV-P5-Vero-P10 virions (**B**) were compared. We did not use the S-IV-P5-Vero-P1 virion in this experiment because the P1 stock retained some carryover virions derived from the Vero-ORF3-E *trans*-complementation culture. Viral titers were analyzed by unpaired T-test. ns, P>0.05.

**Figure S6.**
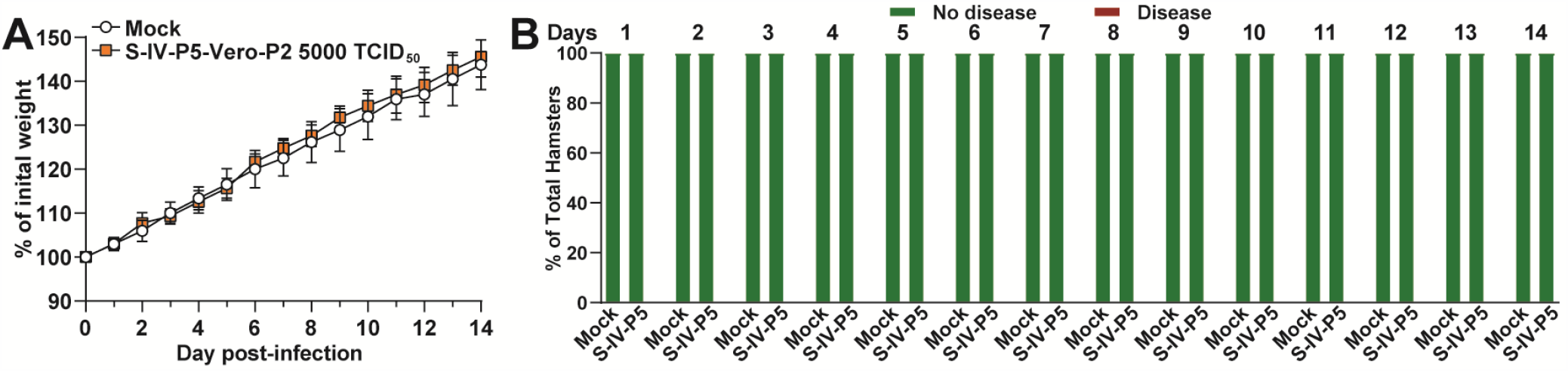
Safety characterization of S-IV-P5-Vero-P2 virion in hamsters. The weight change (**A**) and disease (**B**) of hamsters (n=5) that were intranasally infected with 5,000 TCID_50_ of S-IV-P5-Vero-P2 virion. A high-titer stock of S-IV-P5-Vero-P2 virion used for this experiment was prepared by amplifying the virion on Vero-ORF3-E cells.

**Figure S7.**
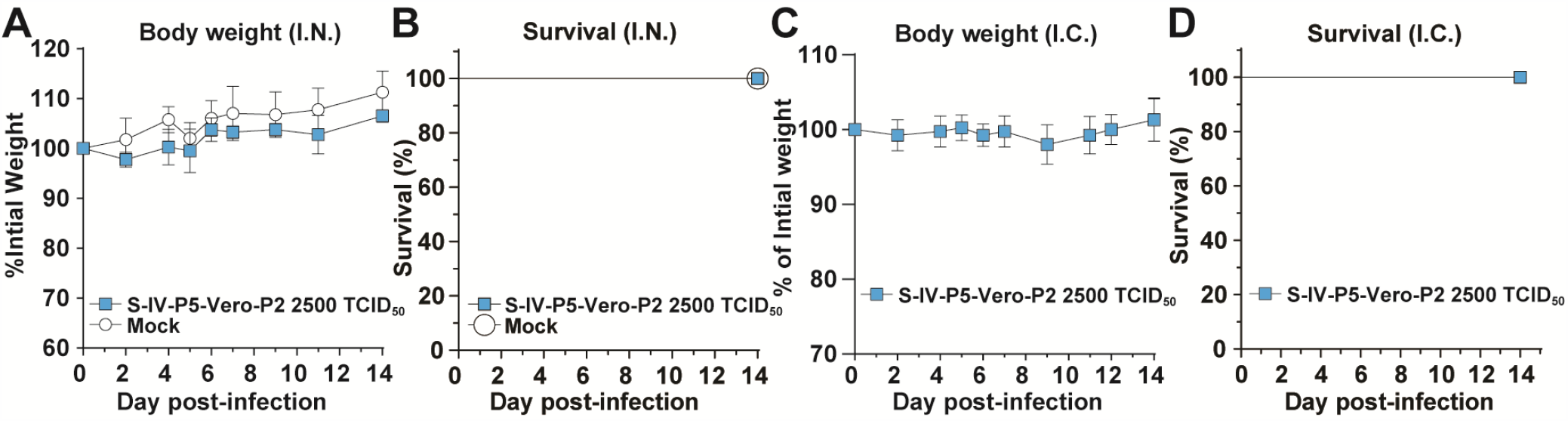
Safety analysis of S-IV-P5-Vero-P2 virion in K18-hACE transgenic mice. Seven- to nine-week-old K18-hACE2 mice were inoculated with S-IV-P5-Vero-P2 virion via the intranasal (I.N.) or intracranial (I.C.) route. Mouse body weight and survival were monitored for 14 days. (**A**) Mouse weight loss after I.N. infection. Mice were infected with 2,500 TCID_50_ of S-IV-P5-Vero-P2 virion (n=4) or PBS mock (n=4) via the I.N. route. The mean ± standard deviations are indicated. (**B**) Mouse survival after I.N. infection. (**C**) Mouse weight loss after I.C. infection. Mouse were inoculated with 500 TCID50 of S-IV-P5-Vero-P2 virion (n=4) via the I.C. route. The mean ± standard deviations are indicated. (**D**) Mouse survival after I.C. infection. A high titer stock of S-IV-P5-Vero-P2 virion used for this experiment was prepared by amplifying the virion on Vero-ORF3-E cells.

**Table S1.** Comparison of neutralization titers between ΔORF3-E mNG virion and PRNT assays.

| Serum ID | ΔORF3-E virion-NT<sub>50</sub> | PRNT<sub>50</sub> |
| --- | --- | --- |
| 1 | <20 | <20 |
| 2 | <20 | <20 |
| 3 | 59 | 80 |
| 4 | 81 | 80 |
| 5 | 169 | 160 |
| 6 | 225 | 200 |
| 7 | 274 | 320 |
| 8 | 353 | 320 |
| 9 | 370 | 320 |
| 10 | 392 | 320 |
| 11 | 394 | 400 |
| 12 | 568 | 320 |
| 13 | 585 | 800 |
| 14 | 666 | 400 |
| 15 | 677 | 640 |
| 16 | 744 | 320 |
| 17 | 909 | 800 |
| 18 | 925 | 640 |
| 19 | 1196 | 800 |
| 20 | 1789 | 1600 |

**Table S2.**
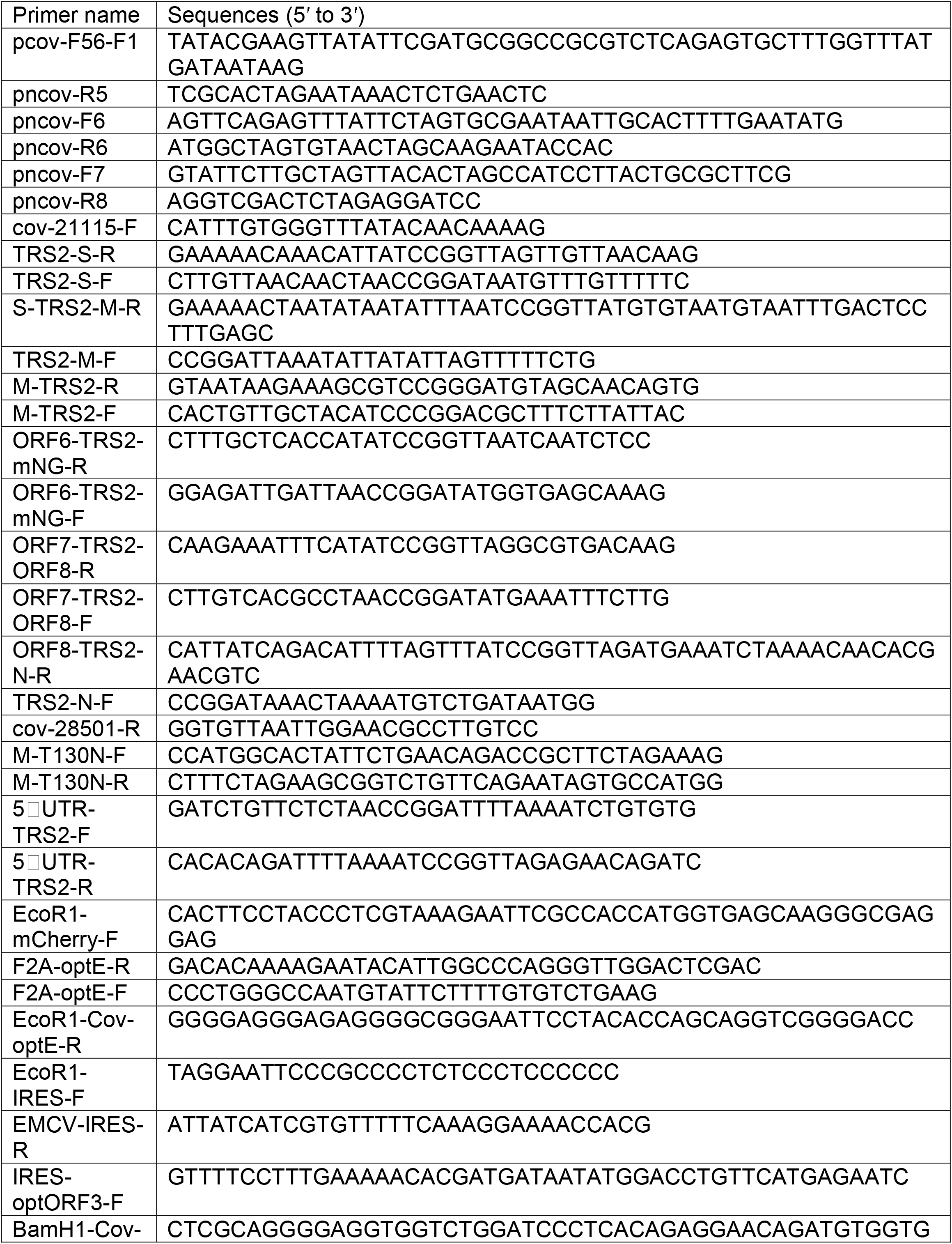
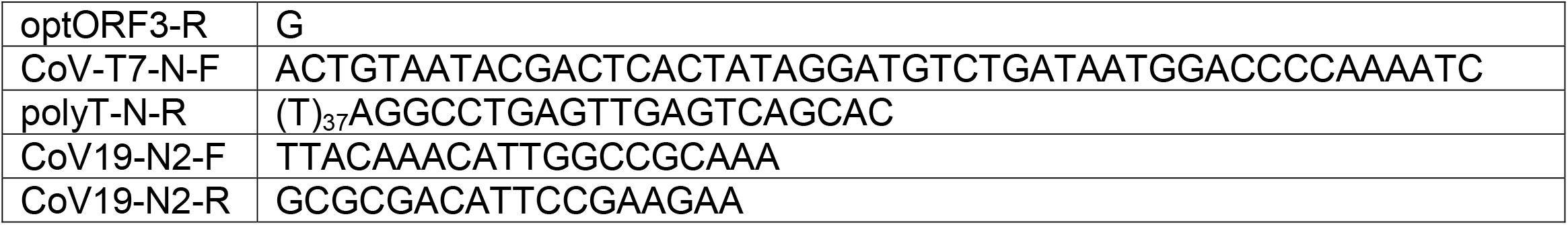
Primers for plasmids construction and RT-PCR analysis.

